# DensiTree 3: Making Sense of Pairs of Sets of Trees

**DOI:** 10.64898/2026.09.28.755190

**Authors:** Remco R. Bouckaert

**Affiliations:** School of Computer Science, University of Auckland Auckland, New Zealand

## Abstract

**Motivation:** Tree sets appear naturally when performing model based phylogenetic analyses. When tracking host-parasite co-evolution, testing sensitivity of different model assumptions, or even just checking convergence of individual runs of the same analysis, the question arises: are a given pair of tree sets the same? And if they are not the same, how do they differ?

**Results:** We provide DensiTree 3, an interactive tool for inspecting pairs of sets of trees that allow visualising both tree sets using a single panel. Incorporation of CCDs provide better summary trees, taxon orderings and extra functionality like rogue detection, making it easier to interpret tree samples. Furthermore, a clade comparison graph allows inspection of differences in clade support and clade heights linked to the double-DensiTree panel. Together, these visualisations of tree sets make it easy to identify potential differences in the tree sets and interactively inspect these differences.

Furthermore, recent advances in conditional clade distributions (CCDs) to represent tree distributions have been incorporated in DensiTree 3, and the most visible part of the user interface has been updated for a more modern look.

**Availability:** DensiTree 3 is freely available from https://cs.auckland.ac.nz/∼remco/DensiTree/. The program is licensed under GPL and source code is available via the link above.

## 1 Introduction

Tree sets are ubiquitous in model based phylogenetic analysis. Co-evolution of host and parasite naturally lead to having two tree sets available representing the evolution of both species, and a desire to find congruence between-the two sets. Fahrenholz’s rule (Fahrenholz [1913]) states that a parasite phylogeny mirrors its host phylogeny, which when performing model based analysis that capture uncertainty in phylogenies naturally leads to comparing pairs of tree sets. Another situation where multiple tree sets arise is when performing an analysis on the same tree set but making different model assumptions, like for example a strict clock versus a relaxed clock. If it turns out the model assumptions do not have an impact on the tree set, and thus on questions of interest, computational expensive and tedious further model selection analyses like path sampling (Baele et al. [2012]) or nested sampling (Russel et al. [2019]) can be skipped. The most common situation where multiple tree sets arise is when performing multiple runs of the same analysis to check that these analyses converge to the same distribution, which is recommended for example when performing Markov chain Monte Carlo (MCMC) analyses (Drummond and Bouckaert [2015]). Observing the same posteriors provides some comfort that the MCMC algorithm is most likely not stuck in a local optimum. In all of the mentioned scenarios there is a need to compare pairs of tree sets.

Tree sets can be compressed to summary trees (Bryant [2003]) and then these summary trees can be compared. Attempts to make sense of pairs of individual trees led to the development of tanglegrams (Venkatachalam et al. [2010], Scornavacca et al. [2011], Matsen et al. [2016]), graphs that show a tree on the left and the right with roots on the outside of the graph, and and lines in the middle to connect the labels. Taxon labels of both trees are ordered in such a way that the connecting lines look nice, e.g. by minimising the number of crossings. An interactive version is available in Dendroscope (Scornavacca et al. [2011]). Disadvantage of summary trees, apart from having to choose one of the many methods (Bryant [2003]), is that topological uncertainty is hard to represent. Consider a tree set to be a distribution over trees, as is common when performing a model based (Bayesian or maximum likelihood) analysis. Then, two tree sets can represent very similar topology support, but the method for summarising the set can flip in favour of one topology for the first set, and another for the second set^1^. Such small difference is magnified in a tanglegram, and shows up as major discordance where there need not be much difference in the underlying tree distributions.

Visualisation of complete sets of trees is less commonly supported by phylogenetic tree visualisation tools. Multi dimensional scaling (MDS) in a space of Robinson Foulds or subtree prune regraft (SPR) distances has been used to visualise single tree sets (Whidden and Matsen IV [2015]), and plotting multiple MDS panels on difference tree sets allows comparison of multiple trees, though it is hard to determine which part of a tree set is the same and where tree sets differ. Furthermore, as far as we are aware, no interactive tools are available to navigate multiple sets of MDS plots.

DensiTree (Bouckaert [2010], Bouckaert and Heled [2014]) takes a different approach and shows all trees in the tree set at the same time, drawing them using transparency, so areas where many trees agree with each other are more densely coloured. This allows visualisation of uncertainty in clade support and as well as internal node heights. Comparing tree sets through DensiTree cannot be done automatically, but requires creating independent images, making sure taxa are in the same order, then using graphic image manipulation tools to mirror and draw them together as for example in Fig. 1 of Lerner et al. [2011], Fig. 2 of McCormack et al. [2012] or Fig. 4 of Degli Esposti et al. [2021].

**Figure 1:**
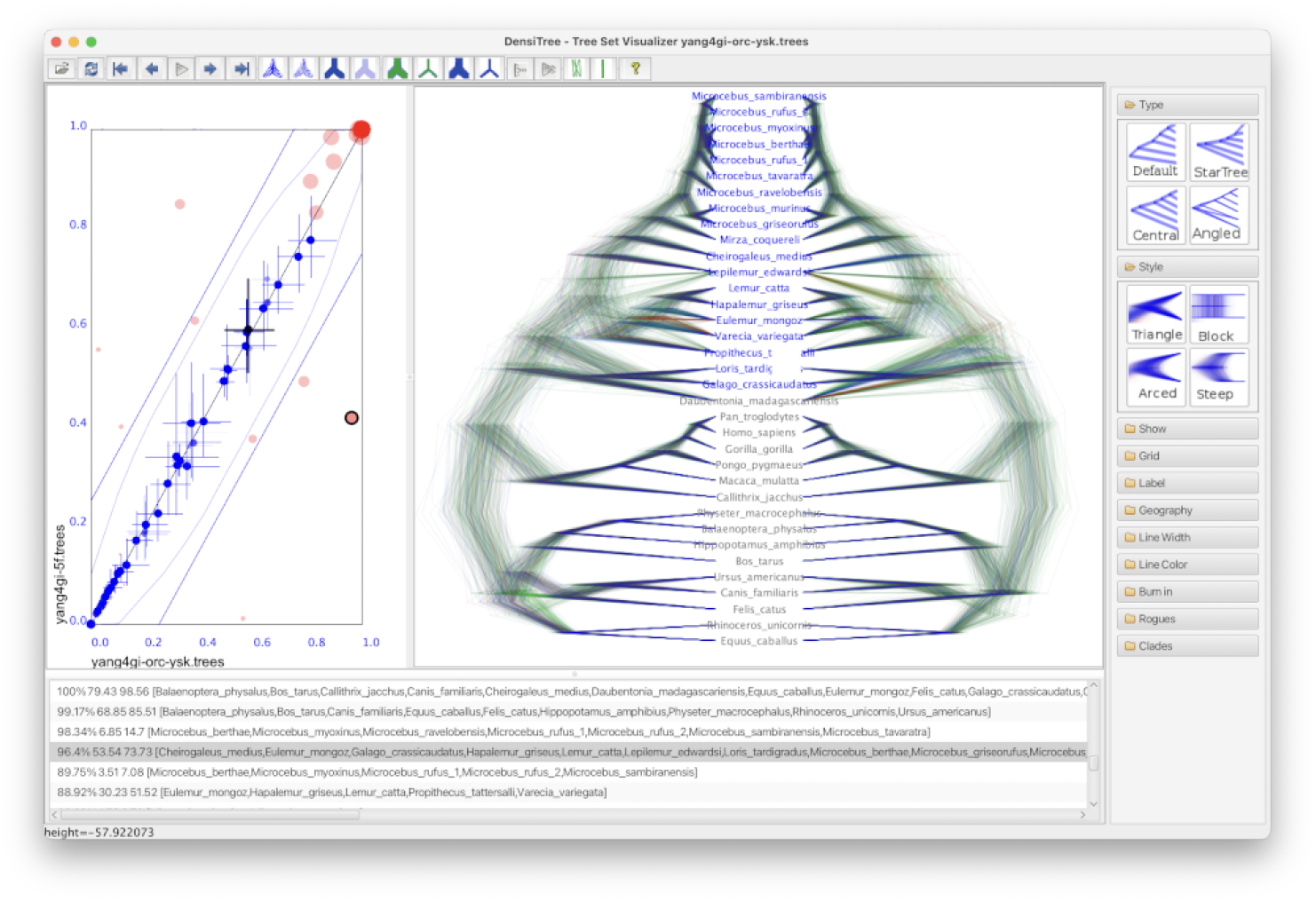
Screen shot of DensiTree 3 showing how two sets of trees can be inspected at the same time and clades of the sets can be compared in the comparison panel on the left hand side.

**Figure 2:**
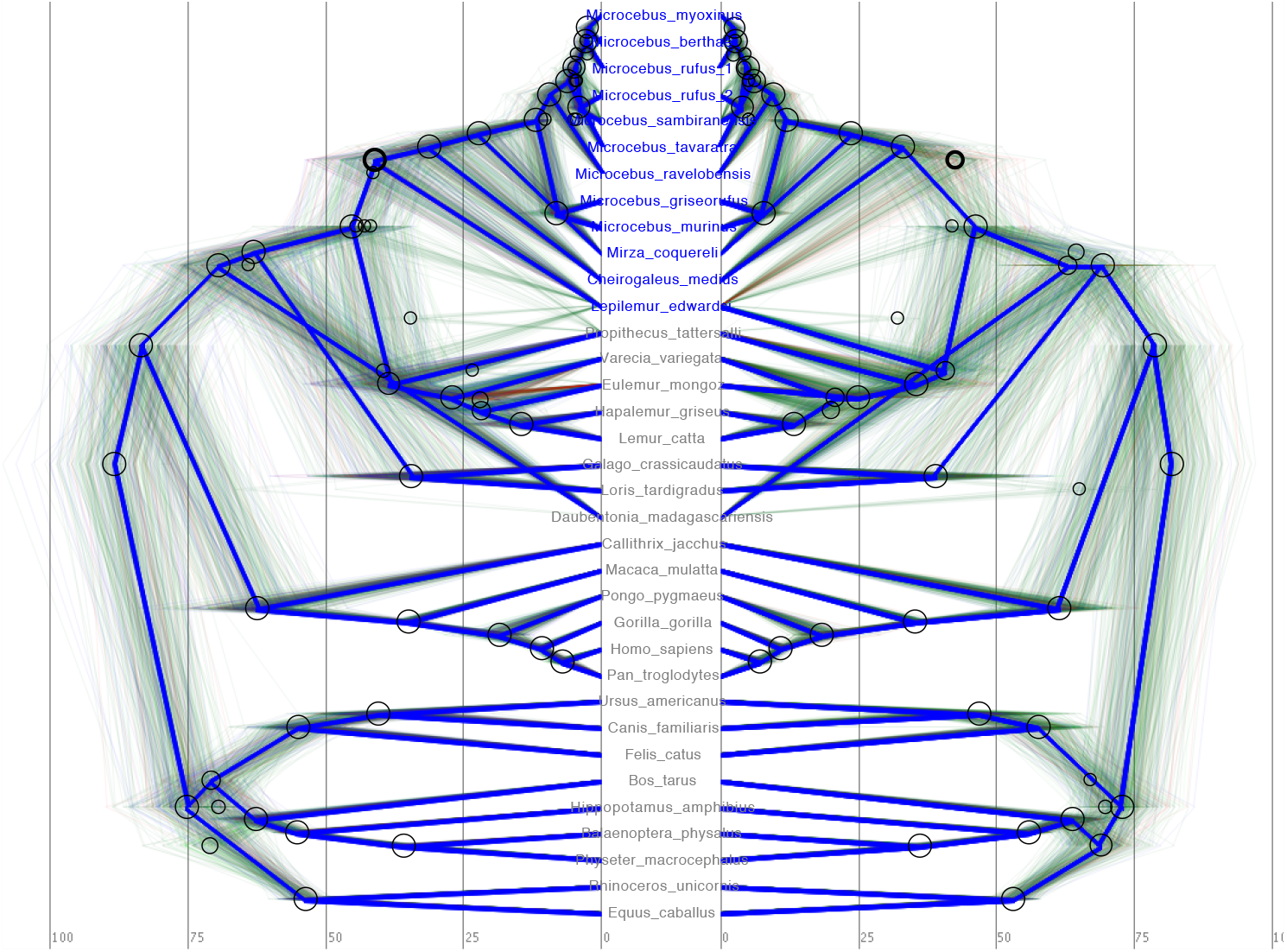
DensiTree 3 showing two sets of trees analysed using different models. Dark blue tree are the summary trees, circles in the DensiTrees indicate clades with over 1% support. The clade where Milne-Edwards’ sportive lemur (*Lepilemur Edwardsi*) flips towards the upper clade in the left hand analysis is selected (blue taxon labels and darker circle for the clade), and the lower clade in the right hand analysis. Other noticeable differences are visible in the summary tree at the bottom around 70 million year where a branch flips over and the placement of the aye aye (*Daubentonia madagascariensis*) which differs in these tree sets.

Here, we present an interactive tool to inspect pairs of tree sets by automatically pairing up the taxa in the tree sets and plot differences in clade support and clade height distributions. These two methods of visualising pairs of tree sets are implemented in version 3 of DensiTree. Furthermore, conditional clade distributions (CCDs) (Höhna and Drummond [2012], Larget [2013], Berling et al. [2025], Klawitter et al. [2024]) are implemented and exploited in a number of ways, including root canal calculation, taxon ordering and rogue detection. The user interface got a facelift by replacing Swing components with JavaFX alternatives (Fig. 1). This required substantial refactoring of the code, but preserved most features of previous versions of DensiTree.

DensiTree is open source, licensed under GPL and available from https://cs.auckland.ac.nz/∼remco/DensiTree/. A manual, including a gallery of tree sets, is available from there as well. Source code is available from https://github.com/rbouckaert/DensiTree.

## 2 Approach

Pairs of trees can be visualised by a paired DensiTree panel and a clade comparison panel. Fig 1 shows an example on a data set from Yang and Yoder [2003] analysed as well in Bouckaert et al. [2019]. These two ways of visualisation will be detailed below.

### 2.1 Paired DensiTree panel

When a single tree set is loaded, DensiTree behaves just like its previous version. When loading a ‘mirror tree set’ as well, the DensiTree panel will show both tree sets with labels in the middle and tree sets on both sides (Fig. 2) provided the taxon sets match. There is no need to have two sets of labels, as in a tanglegram, which does so to prevent line crossings in drawing the two trees. After all, in a DensiTree trees will almost always overlap when there is some level of uncertainty in internal node heights or topology

Taxon label order will be determined by the tree set that was first loaded and shown on the left hand side using one of the built-in node ordering algorithms by default. However, the label ordering can be edited by selecting sets of labels and moving them, or through flipping internal nodes in the ‘edit tree’ as was possible in DensiTree 2 (see manual).

The usual features of DensiTree work for both sets: time scale, visualisation of clade features using visual clues (circles for clade support, bars for clade height) as well as numerical values, displaying a ‘root-canal’ summary tree and colouring branches by metadata in the tree set to name just a few.

### 2.2 Clade comparison panel

The clade comparison panel can be shown next to the pairs of DensiTrees and it shows properties of the clades in both sets (Fig. 3).

**Figure 3:**
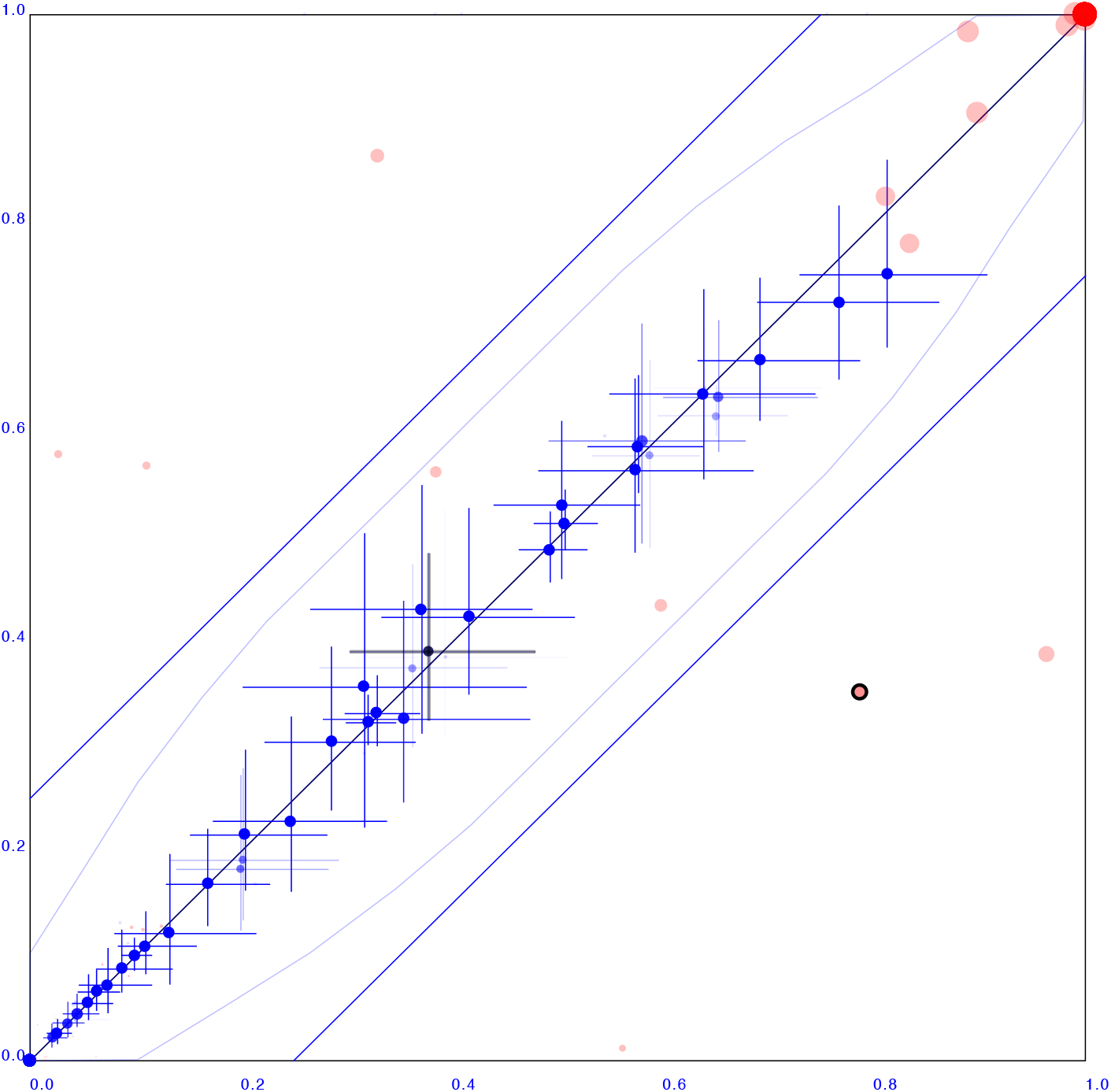
Clade support panel corresponding to the two tree sets shows in Fig. 2. The x-axis represent one tree set, the y-axis the other. Red dots indicate clade support, blue crosses clade height intervals. See text for a more detailed description. The clade where Milne-Edwards’ sportive lemur (*Lepilemur Edwardsi*) flips towards the upper clade in the left hand analysis in Fig. 2 is highlighted both with a dark circle on the clade support entry and by the black instead of blue cross indicating the clade height. Six clades differ more than 25% in support indicated by three red dots above the 25% blue upper line, and three dots below the 25% lower line. They form three pairs corresponding to three topology differences in the summary tree of Fig. 2. See supplementary information for more use cases.

Red dots indicate clade support, running from zero probability (bottom left corner) to 1 (top right corner). When both sets have the same support for a particular clade, the red dot will be on the x=y line, indicated by the diagonal line going from bottom left to top right corner.

The x-axis represent clade values from one set, the y-axis that of mirror set. If both sets are the same, all dots will be on the x=y line (black diagonal line). The two diagonal blue lines indicate there is 25% difference: points within these lines have less than 25% difference, points outside have more, and can be considered as evidence there is substantial difference between the two posteriors.

The upgrade from Swing to JavaFX is visible in the panel on the right hand side and the clade list at the bottom.

Note how the selected clade is highlighted in various places:

- In the clade comparison panel, both the clade support (red dot) and clade height (blue cross) are emphasised in black
- In the tree, taxon labels for taxa in the clade are coloured in blue, the others are greyed out.
- In the clade list panel (at the bottom) the clade is selected as well.

Further, the blue bended lines indicate an envelope of a binomial distribution with 100 trials formed by the 95% upper and lower bounds expected to be seen when two samples are drawn from the same binomial distribution with probability *p* in the range from 0 to 1. Though this is not a formal test that clade set support is estimated differently, red points found outside the envelope indicate probably. Such a test could be based on considering the presence or absence of a clade in the tree set a random binary variable that is binomially distributed. However, since trees in the set are often correlated when obtained by MCMC this approach is hampered by us requiring an effective sample size, which is usually different from the size of the tree set, and can be hard to estimate.

Blue dots indicate the mean of the height of a clade. Both x and y axis are scaled so that the largest tree height (from among both data sets) is at the top right corner. Blue crosses indicate 95% confidence interval of clade height. When both lines of a blue cross intersects the x=y line the clade height distribution can be considered to be compatible. In Fig. 3 all clade height distributions are compatible, though their means can deviate substantially, as shown by the places where the blue dots are some distance away from the x=y line. In contrast, Fig. S1 shows a clade comparison between two sets obtained by two individual MCMC runs on the same model. Blue dots are much closer to the x=y line and all red clade support dots are close to the x=y line as well.

For both red and blue dots, bigger dots mean more clade support. Leaving the clade support dots the same size can result in a large red smear near the lower left corner when there are many clades with low support in both sets. Large support clades are typically less numerous: when there are many clades with substantial support, not much of the trees are available for low support clades.

### 2.3 Selecting clades of interest

Clades can be selected by clicking a clade in one of the two tree sets, or by clicking a clade in the clade comparison panel. When a clade is selected, it is highlighted in the DensiTree by marking the clade with a fatter line than the other clades. In the clade comparison panel, the clade support dot is marked likewise, and the clade height cross is drawn in black instead of blue. Multiple clades can be selected at the same time.

More pairwise statistics like the average difference in mean clade heights, or histograms of differences in branch lengths above a clade can be calculated^2^, but so far have not been found to be as useful in an interactive setting.

Because clades can be selected in the clade comparison panel this makes it very easy to identify which clade differs in both tree sets.

Often, there is uncertainty in tree sets due to short branches and the set contains clades (A,B) and (B,C) with some non-zero level of support. These are often the candidates where the other tree set differs, and where we find significant differences in the summary tree. In fact, all size clades outside the 25% bound in the clade comparison panel in Fig. 1 correspond to differences in the summary tree displayed Fig. 2.

## 3 Conditional clade distributions (CCDs)

CCDs (Höhna and Drummond [2012], Larget [2013]) were introduced to provide a principled, efficient way to model and estimate the probability distribution over phylogenetic tree topologies. One of the properties of CCDs is that a tree topology that is not in the tree set, but all of its constituent splits were observed in other trees, a CCD can still calculate an accurate posterior probability for that tree.

### Summary tree

An extension of the original CCDs (Berling et al. [2025]) called CCD0 was shown to be especially effective in inferring point estimates, which can be used as root canal trees in DensiTrees. The CCD0 based maximum aposteriori (MAP) tree is now the default method for inferring summary trees in DensiTree, though it is still possible to provide a summary tree by hand in Newick format.

### Taxon ordering

The CCD0 MAP tree can be used to order the taxa by finding an ordering that is consistent with a traversal of the summary tree. At each internal node, a decision has to be made whether to traverse left or right. A heuristic that minimises the average sideways movement (i.e. vertical branch lengths when drawning the tree from root on the left to taxa on the right and using block trees) appears particularly effective.

### Rogue detection and removal

CCDs allow for the detection of rogue taxa (Klawitter et al. [2024]), which are taxa that greatly reduce the visibility of the underlying structure of a tree distribution. The biological origin of rogues in a phylogenetic posterior include very short branches, long branch attraction, model misspecification, and lack of informative sites in the alignment. By removing the rogue taxa from the tree set a much clearer picture can emerge, which shows up as a DensiTree with much fewer crossing lines. Rogue taxa are identified by comparing the entropy of the tree set based on a CCD of the full tree set with the entropy of CCDs when taxa are removed (Fig S7). This feature is implemented in DensiTree, and allows for removal of one or more rogue taxa via the ‘rogues’ panel on the left hand side of DensiTree. This is a two step process: first, taxa are ranked in order of rogueness, then a taxon can be selected and that taxon plus all higher ranked taxa are removed. Ranked taxa are labelled with the amount of entropy that is left after removing the taxon.

## 4 Some demonstration applications

### Single tree

Since we can visualise a complete tree set, we can also visualise a single tree. Fig. S2 shows the result of an experiment where a tree is used to simulate an alignment and a new tree inferred using UPGMA, a distance based clustering algorithm. From the clade comparison panel it is immediately obvious that the heights of the true and UPGMA tree are not in agreement.

### True vs. inferred

The result of repeating the analysis with a BEAST2 (Bouckaert et al. [2019]) analysis is shown in Fig. S3. Unlike UPGMA, BEAST produces a tree sample, and the figure shows clade height estimates are consistent with that of the true tree.

### Gene vs. species tree

StarBeast3 (Douglas et al. [2022]) analyses are multi species coalescent analyses, where gene trees and species trees are sampled explicitly during the MCMC. DensiTree 3 makes it possible to visualise both gene trees and species tree at the same time (Fig. S4).

### Cube and matrix vs original

Cube and more generally matrix representation of tree distributions (Bouckaert [2024]) are methods for parameterising a probability distributions over evolutionary trees using continuous Euclidean space. Conceptually, a multivariate (log) normal distribution is used to represent a distribution over entries in a distance matrix, from which the tree is constructed using single link hierarchical clustering. DensiTree 3 can be used to see the biases in cube and matrix representations of tree distributions. Fig. S5 shows how well a cube and a matrix can represent a RSV2 tree sample with 129 taxa. The cube representation does not contains all trees (unlike the matrix) representation, which results in a bias towards higher clade supports for those clades that *are* represented in the cube. This shows up in the red dots mostly lying above the x=y line in Fig. S5. Blue dots at the bottom of the matrix representation show up due to the fact the matrix representation ignores dated tips, even though the input tree set has them. Fig. S6 shows results from the Yang dataset (as in Fig. 2), where the blue dots at the bottom vanished.

## 5 Technical details

DensiTree 3 can read tree sets in NEXUS format (Maddison et al. [1997]), such as those produced by programs like MrBayes (Huelsenbeck and Ronquist [2001]) and BEAST 1 (Suchard et al. [2018]) and 2 (Bouckaert et al. [2019]), and lists of Newick trees as produced by PHYLIP (Felsenstein [1993]). Most aspects of the tree drawing can be configured, including line width, line colour, intensity, font, background colour, etc. and can be passed as command line options. Images can be exported in BMP, JPG, and PNG bitmap image formats and SVG and PDF vector formats.

DensiTree 3 uses its own built in drawing algorithm that uses three (for red, blue and green) 32-bit integers for each pixel, where images in Java typically only use three 8-bit values. Using the built in graphics libraries would result in tree sets larger than 256 (= 2^8^) trees to become so faint that they would not show up any more.

Swing and JavaFX are two standard user interface libraries for Java applications, and DensiTree was written in Swing at a time that JavaFX was not available. For version 3.2, the most visible part of the Swing components have been converted to JavaFX, giving a more modern look and feel than previous versions originating more than a decade ago. The main panel containing the trees and clade set comparison is still a Swing based panel, but since it only contains canvasses used to draw custom panels, conversion to JavaFX would not result in a significantly different user experience. Combining Swing and JavaFX components does require some care, since the different widgets run in their own threads, and one user interface thread cannot update components managed by the other user interface thread.

DensiTree 3 is written in Java, so any computer that runs a Java runtime version 11 or later should be able to use DensiTree 3. The drawing of trees is performed with multiple threads, so that modern multicore machines are fully utilised. Tree sets with ten thousand trees and hundreds of taxa are manageable with DensiTree 3. Very large tree sets with many taxa may take a few minutes to draw to a large extent due to the computational effort required to find a good taxon ordering). Drawing only consensus trees can speed up the process a bit. DensiTree 3 is licensed under the GNU public license. A manual is available via https://cs.auckland.ac.nz/∼remco/DensiTree/ and contains further details of the user interface.

## 6 Open problems

Ordering taxa that maximise readability of paired DensiTrees requires attention. Currently, the ordering is based on the first tree set loaded using heuristics that seem to work based on anecdotal evidence. Even for a single DensiTree, there is no formal criterion agreed upon that quantifies readability and it is unclear whether criteria like entropy of a DensiTree image or the area of a DensiTree without lines would be suitable metrics. Tackling this as first step, finding good taxon ordering algorithms and generalising to paired DensiTrees all are open to further research.

The paired DensiTree gives mostly a qualitative impression about the differences in tree sets. Automating the detection of statistically significant differences requires development of the statistical machinery to be able to so, and integration into DensiTree to visually report the areas of interest. To be able to develop such statistics probably requires that assumptions need to be made about the source of the tree set, like being the output of an MCMC run, which implies potentially correlations between trees and between clades. A statistical test for host-parasite coevolution based on a pair of summary trees already exists (Legendre et al. [2002]), which might be generalised to pairs of sets of trees. Perhaps it is easier to create a statistical test for the situation of comparing posterior tree sets from independent MCMC runs on the same analysis. If these are valid samples from the posterior, these sets should represent exactly the same tree distributions. And this information can be used to inform the statistical test.

When there is considerable topological uncertainty represented by a tree set, DensiTrees can become quite hard to read due to many topologies being drawn on top of each other. Consequently, any structure in the tree set is blurred out when there is a large amount of uncertainty. With pairs of tree sets, this problem is more than doubled, since apart from finding structure in the individual sets, finding similarity between the pairs of sets is hampered by large amounts of uncertainty as well. The clade comparison panel helps to some extent, and rogue removal contributes as well, but the ultimate goal of navigating (pairs of) tree sets with large amounts of uncertainty remains elusive so far in practice.

Ongoing work in tractable tree distributions, in particular conditional clade distributions (Yang et al. [2025], Klawitter and Drummond [2026]) and matrix representations (Bouckaert [2024]) can be an inspiration for further work on analysing tree posteriors. Since properly characterising a single tree distribution is already a formidable task, pairs of tree posteriors have not been given much attention, but may benefit from this work in the future.

## Software availability

DensiTree 3 is open source, licensed under the GNU public license. Source code is available from https://github.com/rbouckaert/DensiTree. The software can be downloaded from https://github.com/rbouckaert/DensiTree/releases. A manual is available via https://cs.auckland.ac.nz/∼remco/DensiTree/ and contains further details of the user interface. Please report bugs and feature requests to https://github.com/rbouckaert/DensiTree/issues.

## Acknowledgement

Joseph Heled was originally an author for his contribution of previous versions of DensiTree, but unfortunately he passed away. This work was greatly assisted by discussions with Alexei Drummond and other members of the Centre of Computational Evolution at the University of Auckland.

## Supplementary information

**Figure S1:**
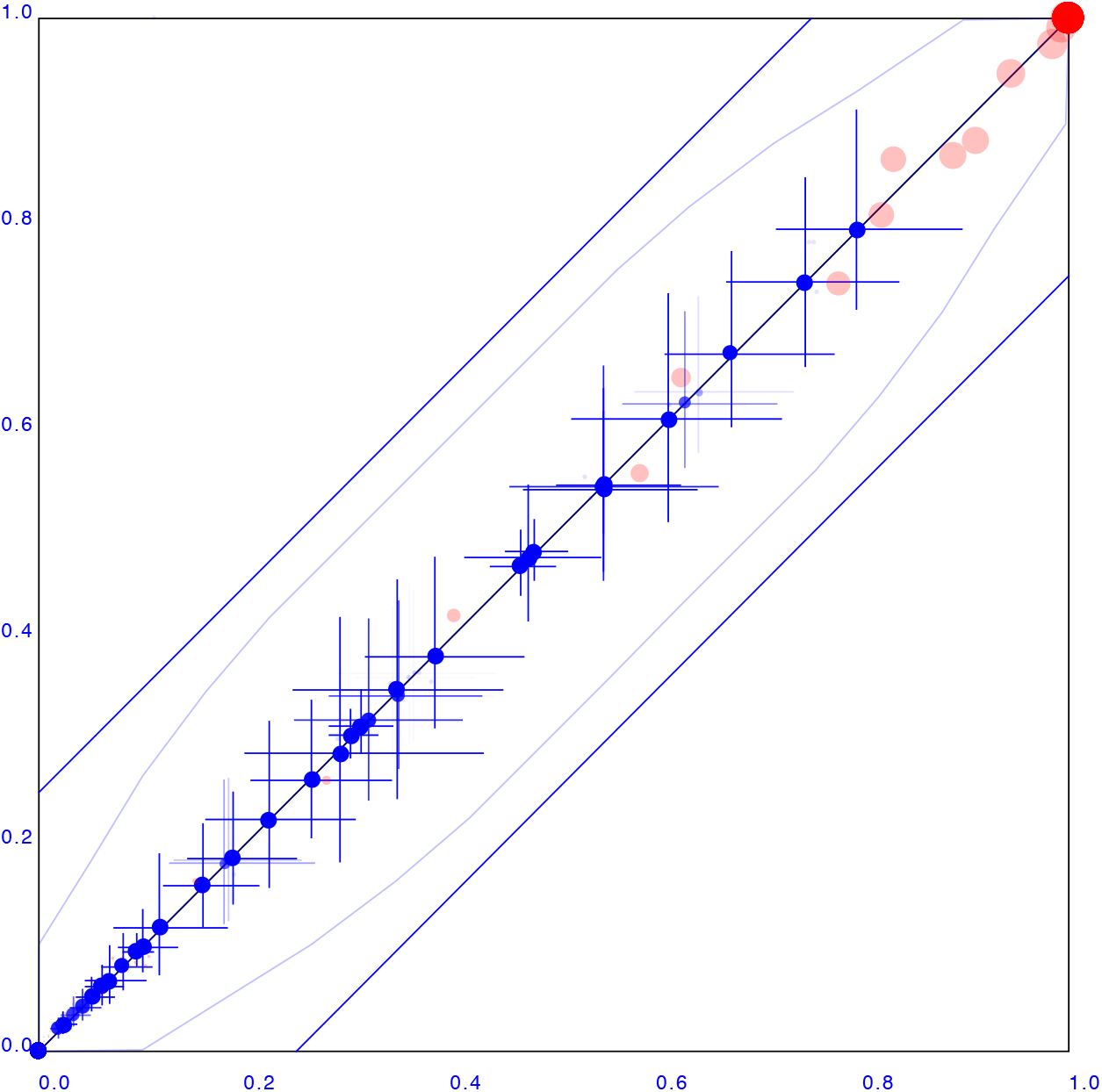
Clade support panel corresponding to the two tree sets from two individual MCMC runs that are fully compatible. In contrast with Fig. 3, the clade support dots in red are much closer to the x=y line, and clade heights shown as blue crosses all look mirrored around the x=y line.

**Figure S2:**
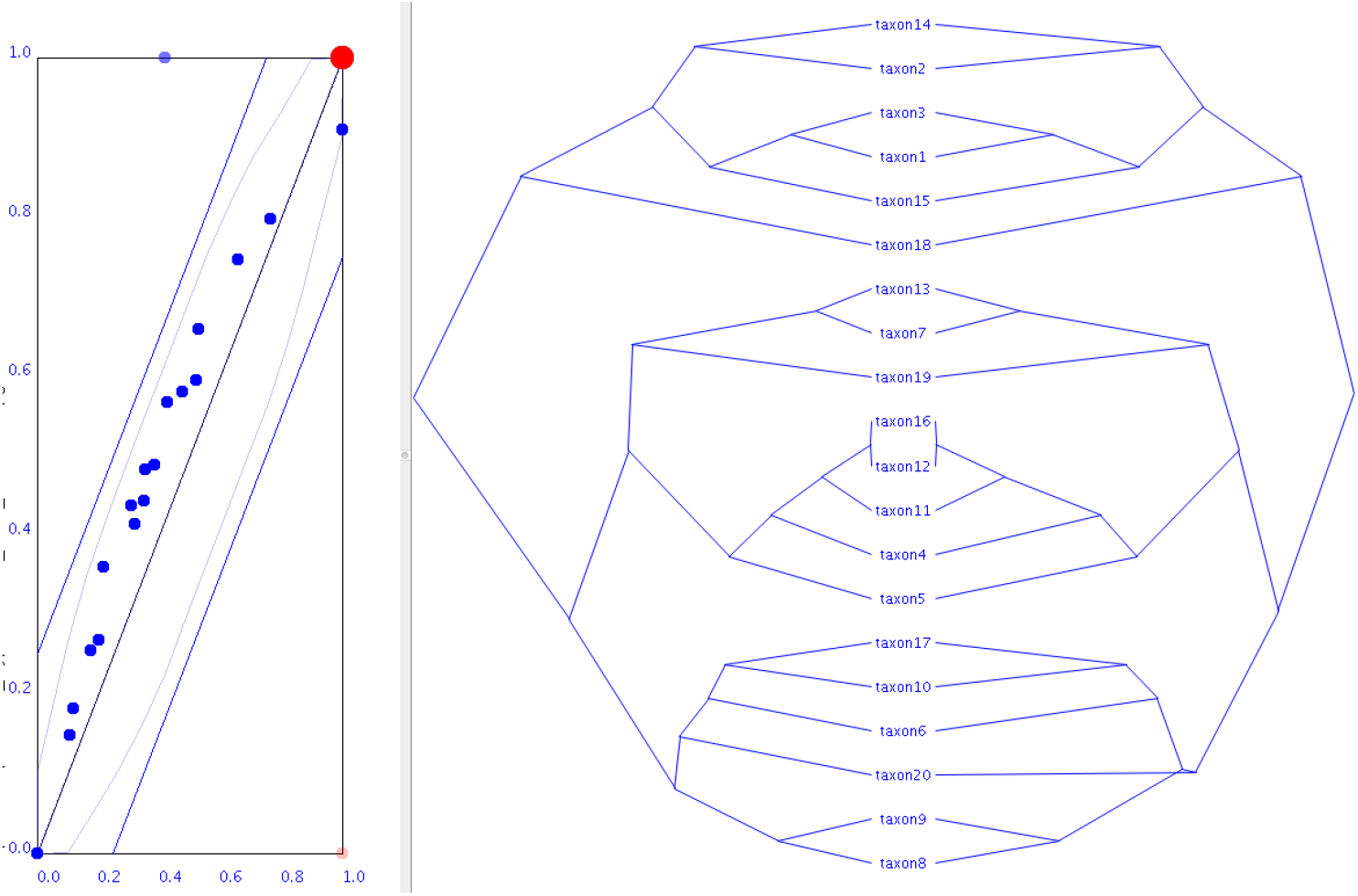
Original tree (left) and UPGMA tree (right) reconstructed from data simulated on the true tree. Note that though the topologies mostly agree (except for a single clade), but clade heights from the original tree differ considerably from that of the UPGMA tree.

**Figure S3:**
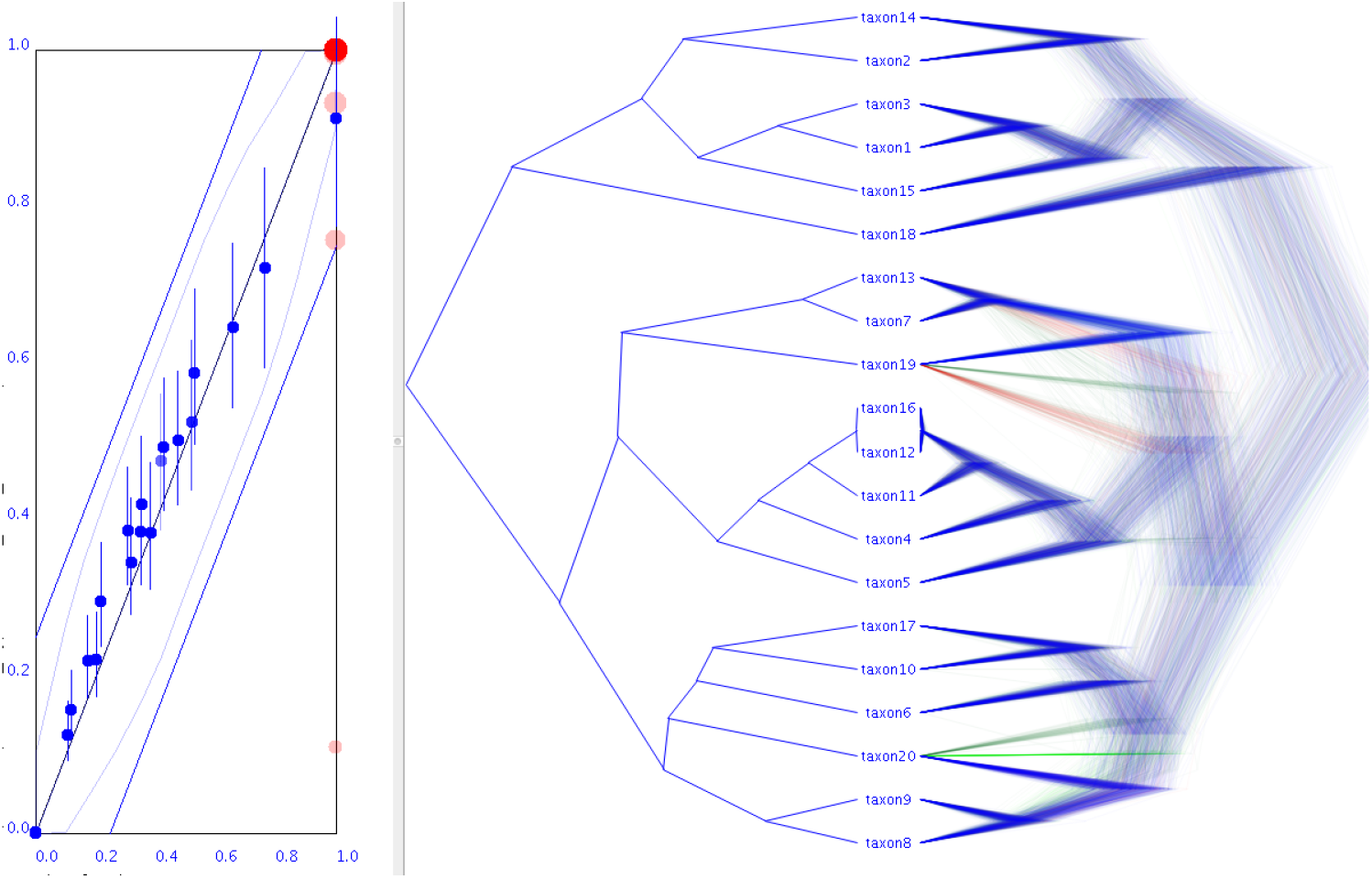
Original tree (left) and BEAST tree (right) reconstructed from data simulated on the true tree. Note that clade height estimates from the BEAST tree set agree with that of the true tree, unlike those of the UPGMA tree in Fig S2. This is shown by the 95%HPD bars in blue consistently containing the true clade height, which we conclude from the fact that the blue bars are all crossing the x=y line.

**Figure S4:**
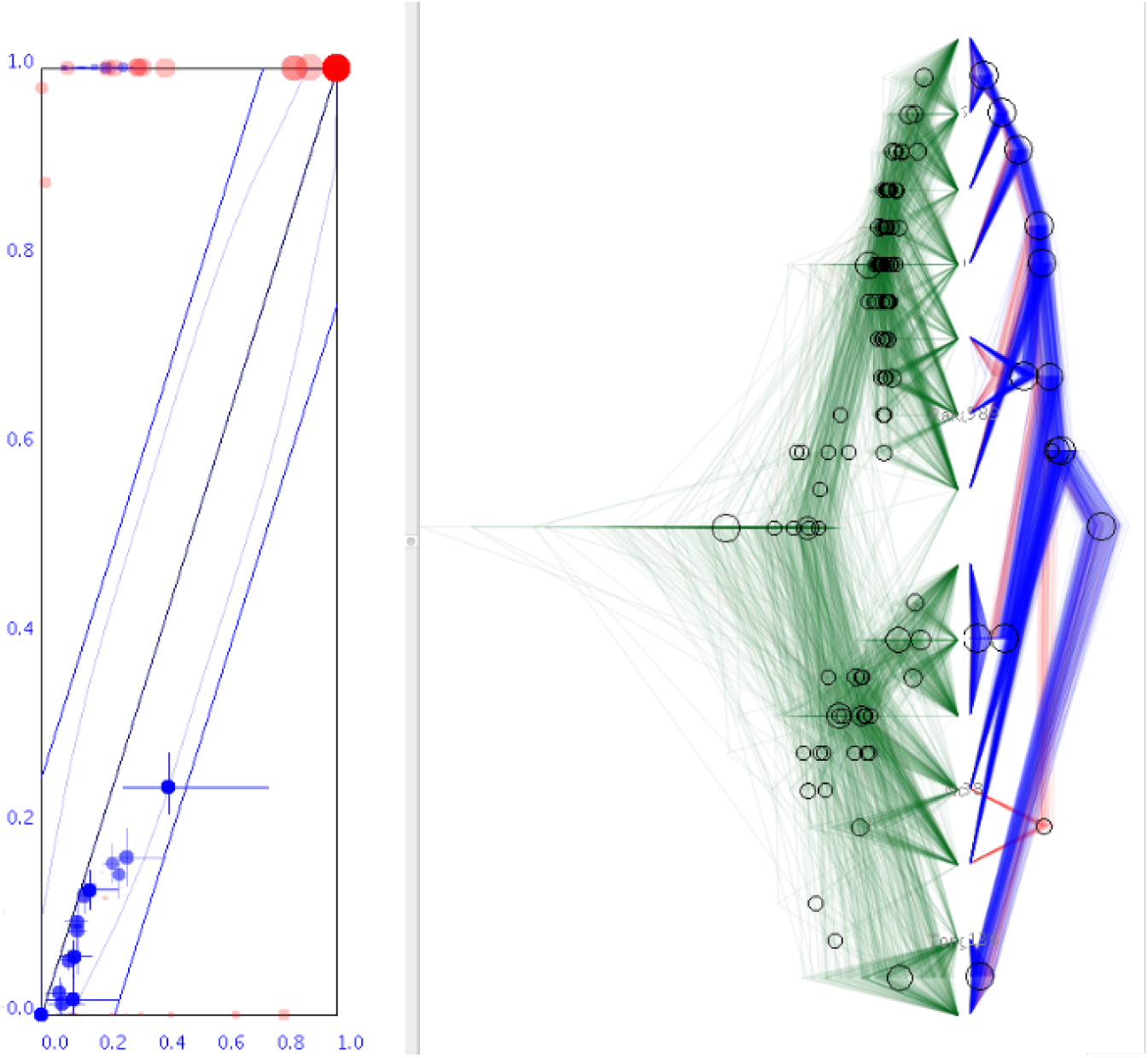
Visualising a multispecies coalescent analysis with StarBeast3 Douglas et al. [2022] with gene trees on the left and species trees on the right. Gene trees have much older internal nodes than the species trees. Clade support differs dramatically as well.

**Figure S5:**
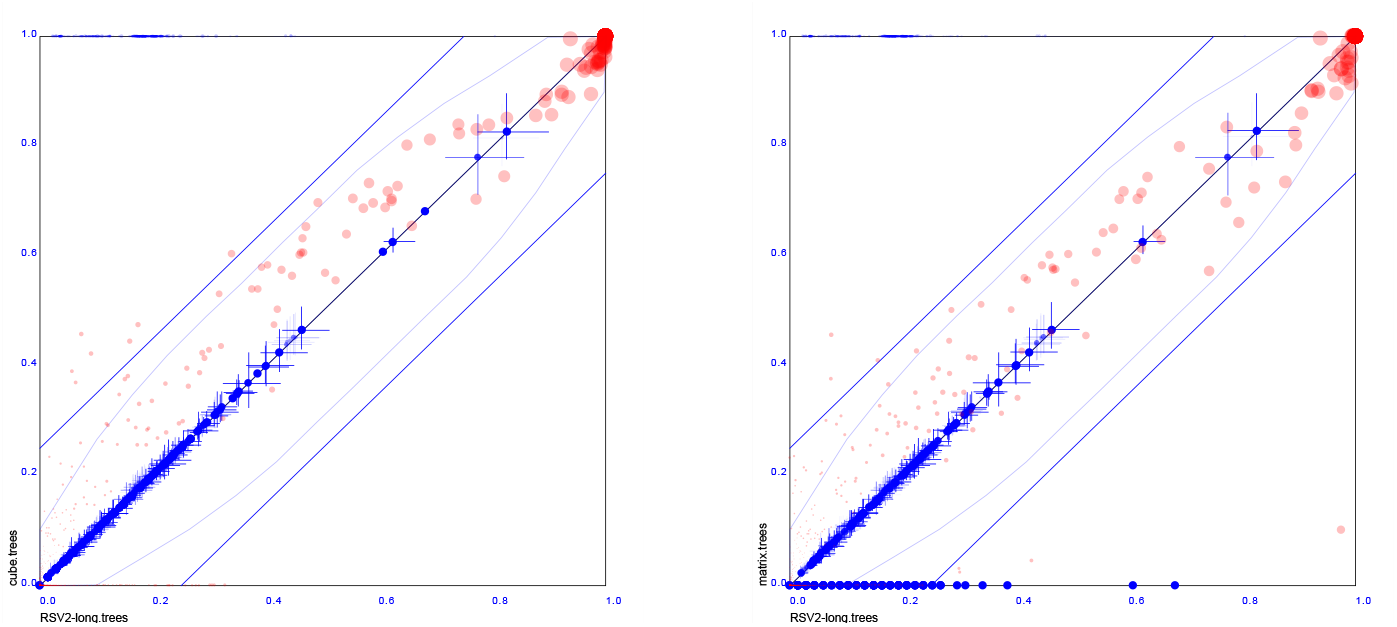
Comparing the RSV2 tree set with sets sampled from a cube representation (left) and from a matrix representation (right). The cube representation over-represents clade support judging from the red dots mainly appearing above the x=y line. The matrix representation is less biased, but still has high variance in clade supports, given many dots are far from the x=y line. Both representations appear to represent clade heights rather well.

**Figure S6:**
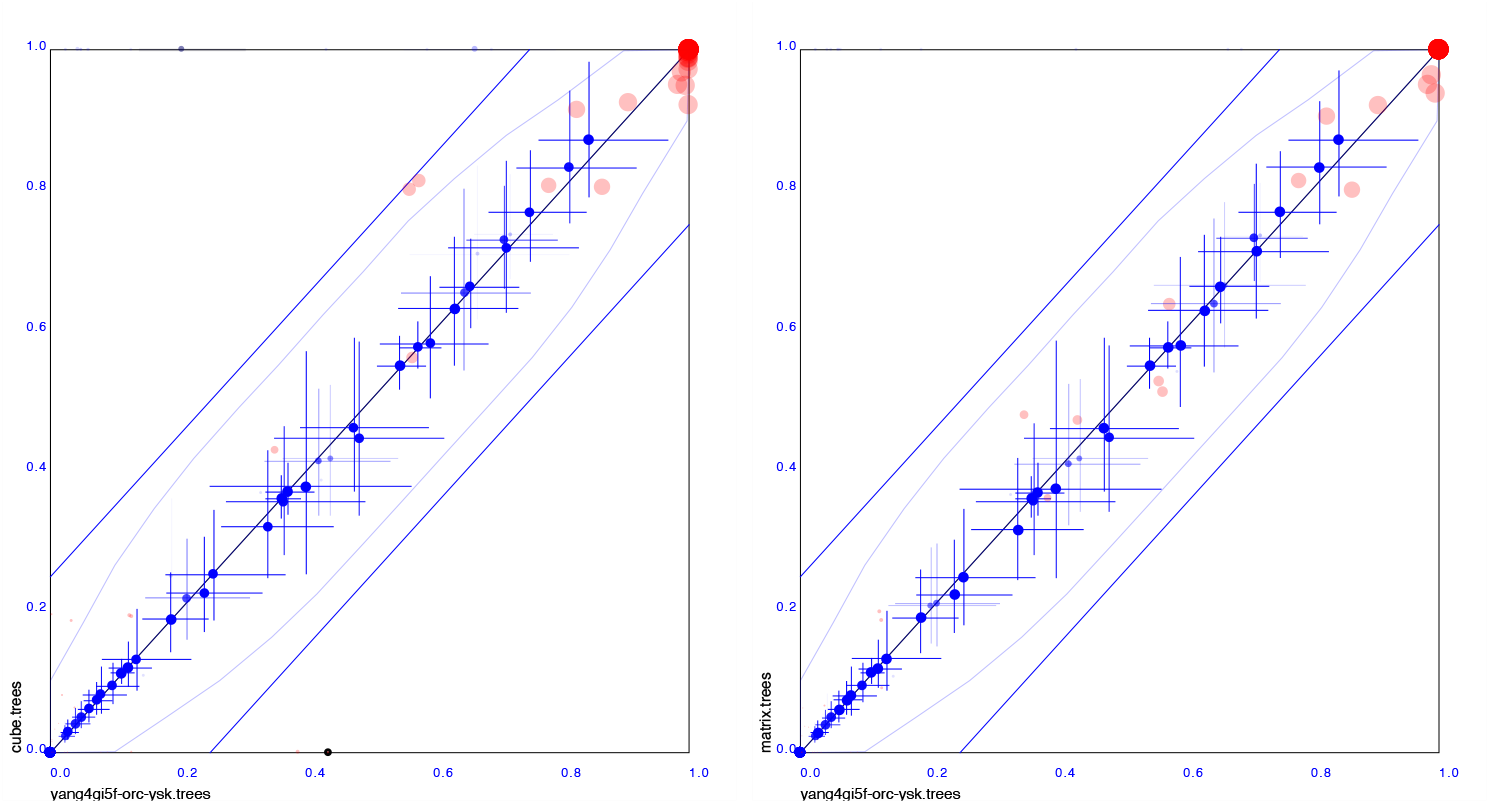
Comparing the Yang tree set in a similar was as Fig. S5.

**Figure S7:**
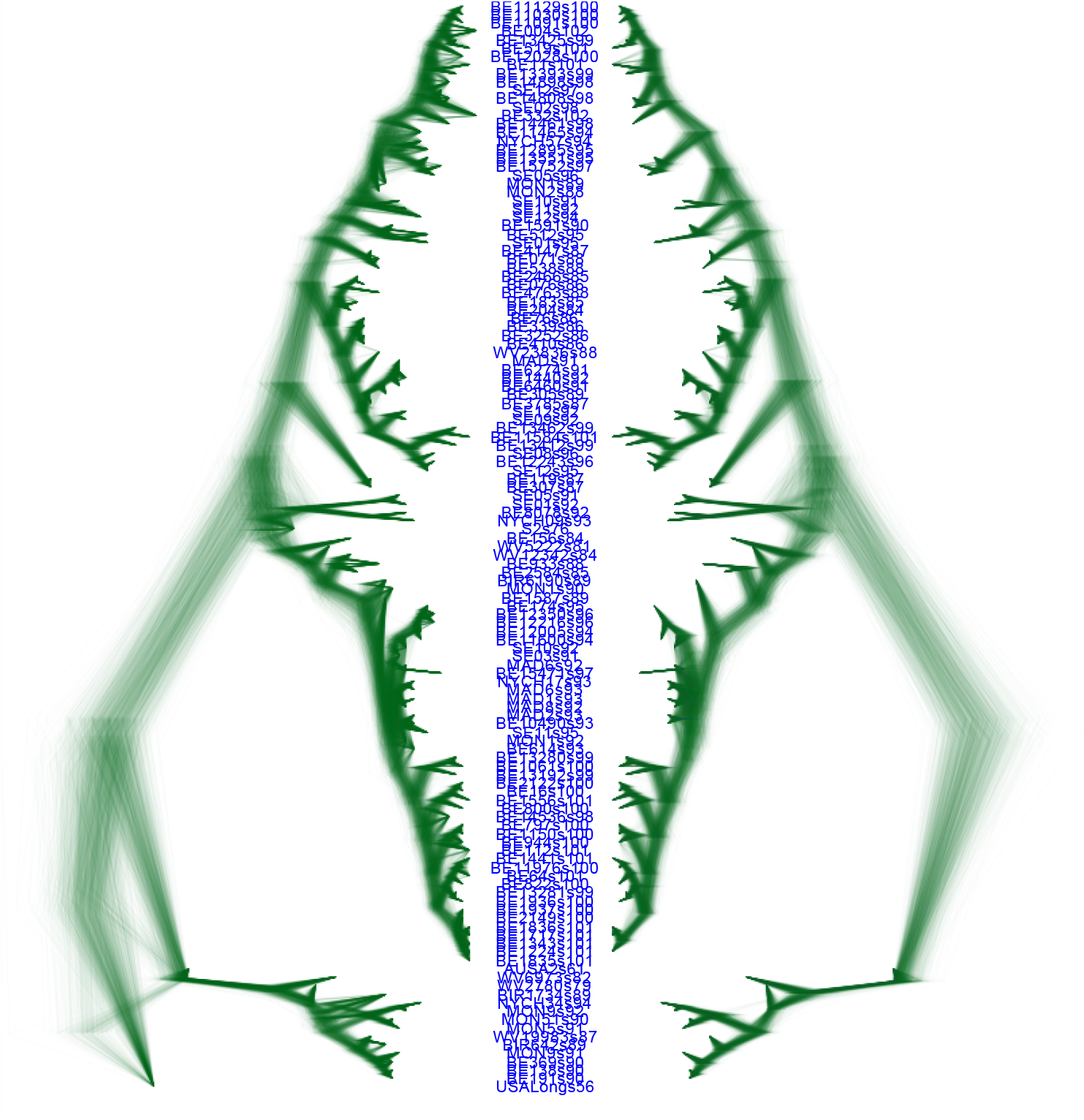
RSV2 analysis with 129 taxa sampled through time, with an entropy of 52.3, so rather diffuse. Removing taxa so that the entropy is reduced to below 10 based on the underlying CCD Klawitter et al. [2024] leaves 91 taxa. These are displayed on the right hand side of the paired DensiTree.

## Footnotes

1 When sampling from the prior, or dealing with identical sequences, this scenario is very common.

2 These are available through the CladeSetComparator tool from the Babel package for BEAST 2 described here: https://www.beast2.org/2020/04/20/comparing-tree-sets.html.

## References

Guy Baele, Philippe Lemey, Trevor Bedford, Andrew Rambaut, Marc A Suchard, and Alexander V Alekseyenko. Improving the accuracy of demographic and molecular clock model comparison while accommodating phylogenetic uncertainty. Molecular biology and evolution, 29(9):2157–2167, 2012.

Lars Berling, Jonathan Klawitter, Remco Bouckaert, Dong Xie, Alex Gavryushkin, and Alexei J Drummond. Accurate Bayesian phylogenetic point estimation using a tree distribution parameterized by clade probabilities. PLOS Computational Biology, 21(2):e1012789, 2025. doi: 10.1371/journal.pcbi.1012789.

Remco Bouckaert, Timothy G Vaughan, Joëlle Barido-Sottani, Sebastián Duchêne, Mathieu Fourment, Alexandra Gavryushkina, Joseph Heled, Graham Jones, Denise Kü hnert, Nicola De Maio, et al. BEAST 2.5: An advanced software platform for Bayesian evolutionary analysis. PLoS computational biology, 15(4):e1006650, 2019.

Remco R Bouckaert. DensiTree: making sense of sets of phylogenetic trees. Bioinformatics, 26(10):1372–1373, 2010.

Remco R Bouckaert. Variational bayesian phylogenies through matrix representation of tree space. PeerJ, 12:e17276, 2024.

Remco R Bouckaert and Joseph Heled. DensiTree 2: Seeing trees through the forest. BioRxiv, page 012401, 2014.

David Bryant. A classification of consensus methods for phylogenetics. DIMACS series in discrete mathematics and theoretical computer science, 61:163–184, 2003.

Mauro Degli Esposti, Ana Moya-Beltrán, Raquel Quatrini, and Lars Hederstedt. Respiratory heme A-containing oxidases originated in the ancestors of iron-oxidizing bacteria. Frontiers in Microbiology, 12:1465, 2021.

Jordan Douglas, Cinthy L Jiménez-Silva, and Remco Bouckaert. Starbeast3: adaptive parallelized Bayesian inference under the multispecies coalescent. Systematic Biology, 71(4):901–916, 2022.

Alexei J Drummond and Remco R Bouckaert. Bayesian evolutionary analysis with BEAST. Cambridge University Press, 2015.

H Fahrenholz. Ectoparasiten und abstammungslehre. Zoologischer Anzeiger, 41(8):371–374, 1913.

Joseph Felsenstein. PHYLIP (phylogeny inference package), version 3.5 c. Joseph Felsenstein., 1993.

John P Huelsenbeck and Fredrik Ronquist. MrBayes: Bayesian inference of phylogenetic trees. Bioinformatics, 17(8):754–755, 2001.

Sebastian Höhna and Alexei J. Drummond. Guided tree topology proposals for Bayesian phylogenetic inference. Systematic Biology, 61(1):1–11, 2012. doi: 10.1093/sysbio/syr074.

Jonathan Klawitter and Alexei J Drummond. Bayesian credible sets for phylogenetic tree topologies with applications to coverage analysis and cross-model comparison. Molecular Biology and Evolution, 43(7):msag141, 2026.

Jonathan Klawitter, Remco R Bouckaert, and Alexei J Drummond. Skeletons in the forest: Using entropy-based rogue detection on Bayesian phylogenetic tree distributions. bioRxiv, pages 2024–09, 2024.

Bret Larget. The estimation of tree posterior probabilities using conditional clade probability distributions. Systematic Biology, 62(4):501–511, 2013. doi: 10.1093/sysbio/syt014.

Pierre Legendre, Yves Desdevises, and Eric Bazin. A statistical test for host–parasite coevolution. Systematic biology, 51(2):217–234, 2002.

Heather RL Lerner, Matthias Meyer, Helen F James, Michael Hofreiter, and Robert C Fleischer. Multilocus resolution of phylogeny and timescale in the extant adaptive radiation of Hawaiian honeycreepers. Current Biology, 21(21):1838–1844, 2011.

David R Maddison, David L Swofford, and Wayne P Maddison. NEXUS: an extensible file format for systematic information. Systematic biology, 46(4):590–621, 1997.

Frederick A Matsen, Sara C Billey, Arnold Kas, and Matjaž Konvalinka. Tanglegrams: a reduction tool for mathematical phylogenetics. IEEE/ACM transactions on computational biology and bioinformatics, 15 (1):343–349, 2016.

John E McCormack, Brant C Faircloth, Nicholas G Crawford, Patricia Adair Gowaty, Robb T Brumfield, and Travis C Glenn. Ultraconserved elements are novel phylogenomic markers that resolve placental mammal phylogeny when combined with species-tree analysis. Genome research, 22(4):746–754, 2012.

Patricio Maturana Russel, Brendon J Brewer, Steffen Klaere, and Remco R Bouckaert. Model selection and parameter inference in phylogenetics using nested sampling. Systematic biology, 68(2):219–233, 2019.

Celine Scornavacca, Franziska Zickmann, and Daniel H Huson. Tanglegrams for rooted phylogenetic trees and networks. Bioinformatics, 27(13):i248–i256, 2011.

Marc A Suchard, Philippe Lemey, Guy Baele, Daniel L Ayres, Alexei J Drummond, and Andrew Rambaut. Bayesian phylogenetic and phylodynamic data integration using BEAST 1.10. Virus evolution, 4(1):vey016, 2018.

Balaji Venkatachalam, Jim Apple, Katherine St John, and Daniel Gusfield. Untangling tanglegrams: Comparing trees by their drawings. IEEE/ACM Transactions on Computational Biology and Bioinformatics, 7(4):588–597, 2010.

Chris Whidden and Frederick A Matsen IV. Quantifying MCMC exploration of phylogenetic tree space. Systematic biology, 64(3):472–491, 2015.

Ziheng Yang and Anne D Yoder. Comparison of likelihood and Bayesian methods for estimating divergence times using multiple gene loci and calibration points, with application to a radiation of cute-looking mouse lemur species. Systematic biology, 52(5):705–716, 2003.

Zijing Yang, Jonathan Klawitter, Remco Bouckaert, and Alexei J Drummond. Improved phylogenetic posterior estimation through regularised conditional clade distributions. bioRxiv, pages 2025–12, 2025.

